# A new variant of ASIC2 mediates sodium retention in nephrotic syndrome

**DOI:** 10.1101/2021.02.09.430231

**Authors:** Marc Fila, Ali Sassi, Gaëlle Brideau, Lydie Cheval, Luciana Morla, Pascal Houillier, Christine Walter, Michel Gennaoui, Laure Collignon L, Mathilde Keck, Gabrielle Planelles, Naziha Bakouh, Michel Peuchmaur, Georges Deschênes, Ignacio Anegon, Séverine Remy, Bruno Vogt, Gilles Crambert, Alain Doucet

**Author notes:** Present address: Marc Fila, Pediatric Nephrology Department, Hôpital Arnaud de Villeneuve Institut de Génomique Fonctionnelle UMR9023 CNRS U661 INSERM, Montpellier, France; Ali Sassi, Department of Cellular Physiology and Metabolism, University of Geneva, Geneva, Switzerland; Michel Genaoui, 28B Allée des dauphins, Trois Bassins, France; Laure Collignon, 30 rue Jacques Brel, Achères, France; Mathilde Keck, Université Paris-Saclay, CEA, INRAE, Département Médicaments et Technologies pour la Santé (DMTS), SIMoS, 91191 Gif-sur-Yvette, France. Marc Fila and Ali Sassi contributed equally to this work. Correspondence to:Gilles Crambert, CRC, 15 rue de l’Ecole de Médecine, 75270 Paris cedex 6, France, Fax: (33), Phone: (33).

## Abstract

Idiopathic nephrotic syndrome (INS) is characterized by proteinuria and renal Na retention leading to oedema. This Na retention is usually attributed to epithelial sodium channel (ENaC) activation following plasma aldosterone increase. However, most nephrotic patients show normal aldosterone levels. Using a corticosteroid-clamped rat model of INS (CC-PAN), we showed that the observed electrogenic and amiloride-sensitive Na retention could not be attributed to ENaC. We, then, identified a truncated variant of acid sensing ion channel 2b (ASIC2b) that induced sustained acid-stimulated sodium currents when co-expressed with ASIC2a. Interestingly, CC-PAN nephrotic ASIC2b-null rats did not develop sodium retention. We finally showed that expression of the truncated ASIC2b in kidney was dependent on the presence of albumin in the tubule lumen and activation of ERK in renal cells. Finally, the presence of ASIC2 mRNA was also detected in kidney biopsies from patients with INS but in any of the patients with other renal diseases. We have, therefore, identified a novel variant of ASIC2b responsible for the renal Na retention in the pathological context of INS.

## Introduction

Nephrotic syndrome is defined by the association of massive proteinuria with hypoalbuminemia brought about by dysfunctions of the glomerular filtration barrier. Glomerular dysfunctions stem either from poorly characterized immunological disorders or from genetic alterations of glomerular proteins. Whatever its origin, idiopathic nephrotic syndrome is associated with sodium retention which participates in the formation of ascites and edema (Doucet *et al*, 2007). The site and mechanism of sodium retention have been deciphered using the puromycin aminonucleoside-induced rat model of nephrotic syndrome (PAN rat) that reproduces most biological and clinical signs of the human disease (Frenk *et al*, 1955; Pedraza-Chaverri *et al*, 1990). Sodium retention in PAN rats originates from the aldosterone-sensitive distal nephron (ASDN) and stems from the marked stimulation of the basolateral Na^+^,K^+^-ATPase and of the apical sodium channel ENaC, two molecular targets of aldosterone (Deschenes *et al*, 2001; Ichikawa *et al*, 1983; Lourdel *et al*, 2005).

However, conversely to PAN rats most nephrotic patients display normal volemia and plasma aldosterone levels (Vande Walle *et al*, 1995). This raises the possibility that the site and mechanism of sodium retention in most nephrotic patients might be different from those described in PAN rats. As a matter of fact, when their plasma level of corticosteroids is clamped to basal level, PAN rats still develop nephrotic syndrome and amiloride-sensitive edema and ascites, but the mechanism of sodium retention is different as they display no membrane expression and activity of ENaC in the ASDN (de Seigneux *et al*, 2006; Lourdel *et al.*, 2005). The principal aims of this study were to identify the amiloride-sensitive and ENaC-independent sodium reabsorption pathway responsible for sodium retention in these animals, and its mechanism of induction. The secondary aim was to evaluate whether this alternate sodium reabsorption pathway is present in nephrotic patients.

Results show that sodium retention in corticosteroid-clamped (CC) PAN rats originates from ASDN and is dependent on the expression of a short variant of the acid sensing ion channel (ASIC) 2b regulatory subunit that confers to the transiently active ASIC2a the properties of a long-lasting epithelial sodium channel. Expression of this ASIC2b variant during nephrotic syndrome is secondary to albumin endocytosis and activation of ERK pathway in the ASDN. Expression of ASIC2 was also found in ASDN from a majority of nephrotic patients.

## Results

### Sodium retention originates from collecting ducts of CC-PAN rats but is independent of the epithelial sodium channel ENaC

Measurement of the net transepithelial flux of sodium (J_Na_+) by *in vitro* microperfusion of isolated cortical collecting ducts (CCD) showed that, in contrast to CCDs from control rats in which no J_Na_+ is measurable (Morla *et al*, 2013), CCDs from both PAN and CC-PAN rats displayed significant and similar J_Na_+ (figure 1A). CCD from PAN and CC-PAN rats also displayed a lumen negative transepithelial voltage (in mV ± SE; PAN: −14.1 ± 2.2, n=8; CC-PAN: −12.9 ± 1.9, n=5; NS), indicating that sodium reabsorption is an electrogenic process. As previously reported in PAN rats (Deschenes *et al.*, 2001), luminal addition of amiloride abolished J_Na_+ in CCD from CC-PAN rats (figure 1B) as well as the lumen negative transepithelial voltage (in mV ± SE; Control: −9.9 ± 4.7; Amiloride: 5.9 ± 0.8, n=3; p<0.03). All these properties are consistent with an amiloride-sensitive process mediating sodium reabsorption in CCD from CC-PAN rats. However previous studies concluded to the absence of functional ENaC under these conditions (de Seigneux *et al.*, 2006; Lourdel *et al.*, 2005). Using sodium-depleted (LNa) rats as a model of over expression of ENaC in the CCD (Leviel *et al*, 2010), we searched for properties differentiating ENaC-mediated sodium transport from that in CC-PAN rats. J_Na_+ in CCD from LNa rats was markedly reduced following luminal addition of 300μM ZnCl_2 w_hereas it was slightly increased in CC-PAN rats (figure 1C). Luminal acidification (pH ≈ 6.0) abolished J_Na_+ in LNa rats but had no significant effect in CC-PAN rats (figure 1D). Along with pieces of evidence previously reported (de Seigneux *et al.*, 2006; Lourdel *et al.*, 2005), these findings demonstrate that sodium retention in CC-PAN rats originates from the ASDN and stems from the activation of an electrogenic, amiloride-sensitive, Zn- and pH-insensitive transport pathway independent of ENaC.

**Figure 1.**
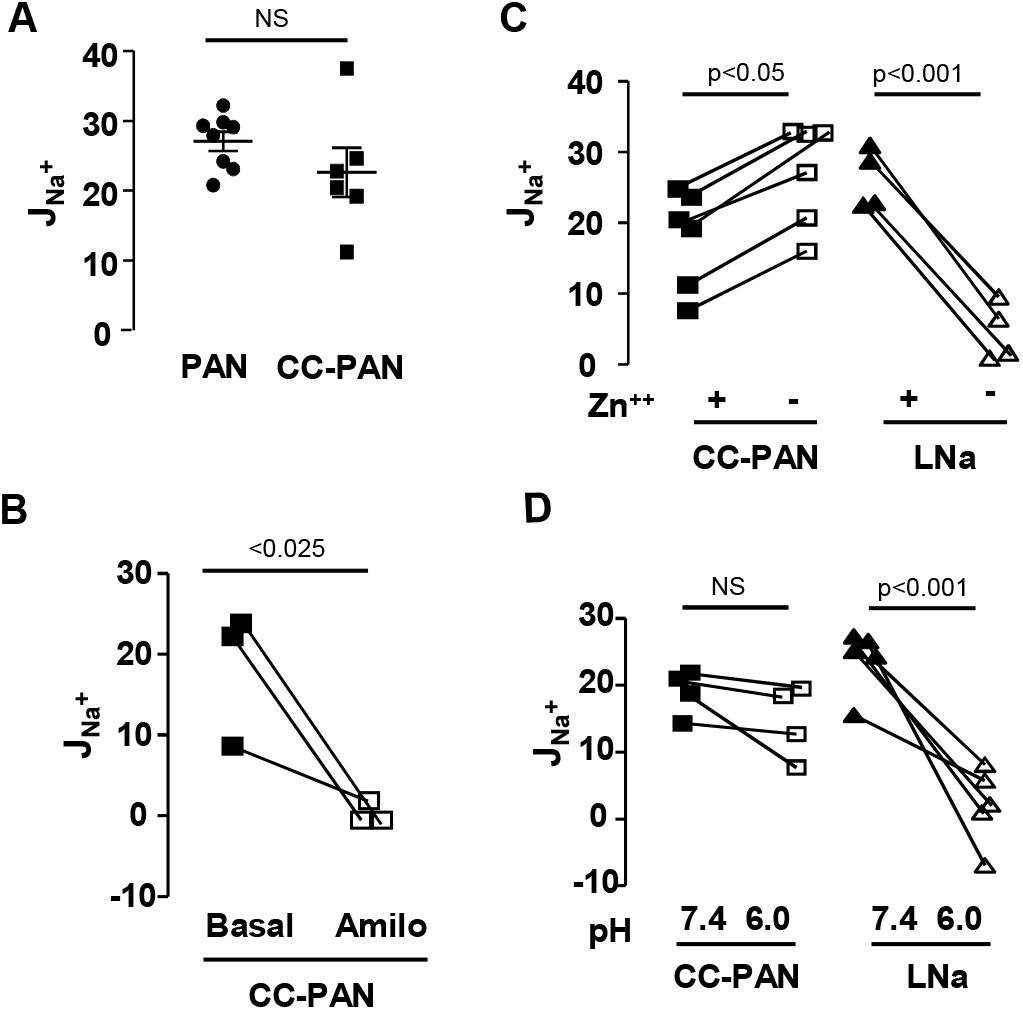
Sodium flux (J_Na_+) measured in microperfused CCDs *in vitro*. **A.** J_Na_+ in CCDs from PAN nephrotic and corticosteroid-clamped PAN nephrotic rats (PAN and CC-PAN). Each value represents a rat. **B.** J_Na_+ was measured in CCDs from CC-PAN nephrotic rats before and after luminal addition of 10μM amiloride. Each value represents a rat. **C.** J_Na_+ in CCDs from CC-PAN rats and rats fed a sodium depleted diet for 14 days (LNa) before and after luminal addition of 300μM ZnCl_2._ Values are means ± SE from 4 LNa and 3 CC-PAN animals. **D.** J_Na_+ in CCDs from CC-PAN and LNa rats was measured before and after acidification of the luminal fluid at pH 6.0. Values are means ± SE from 5 LNa and 4 CC-PAN animals.

### ASIC2 is responsible for J_Na_+ and sodium retention in CC-PAN rats

We therefore looked for the expression in the CCD of CC-PAN rats of amiloride-sensitive channels of the ENaC/degenerin family, which includes the subunits of ENaC and of acid sensitive ion channels (ASICs) (Kellenberger & Schild, 2015). Alike ENaC which is a trimer of α, β and γ subunits, ASICs are homo- or hetero-trimers of ASIC1-5 channel isoforms. Besides the three subunits of ENaC, RT-qPCR, revealed the mRNA expression of ASIC1a, ASIC2a and ASIC2b whereas ASIC1b and ASIC3-5 were not detected. ASIC1a and ASIC2a can constitute functional channels by themselves whereas ASIC2b cannot and is therefore considered as a regulatory subunit that associates with conductive isoforms and modifies their properties (Kellenberger & Schild, 2002). As compared with CC-control rats, the levels of βENaC, γENaC, ASIC1a and ASIC2a mRNAs were unchanged in CCD from CC-PAN rats and that of αENaC was reduced by 40%. In contrast, the expression level of ASIC2b mRNA was increased 2-fold in CCD from CC-PAN rat (figure 2A). This suggests that ASIC2b in association with ASIC1a and/or ASIC2a may participate in J_Na_+ in CCD and in sodium retention in CC-PAN rats. This hypothesis was confirmed by the finding that genetic deletion of ASIC2b (see Supplementary Data) abolished the stimulation of J_Na_+ in CCD and sodium retention in CC-PAN rats, and reduced by over half the volume of ascites without altering proteinuria (Figure 2B-D). The residual volume of ascites observed in CC-PAN ASIC2b^−/−^ rats is likely accounted for by the increased permeability of peritoneal capillaries described in PAN rats (Udwan *et al*, 2016).

**Figure 2.**
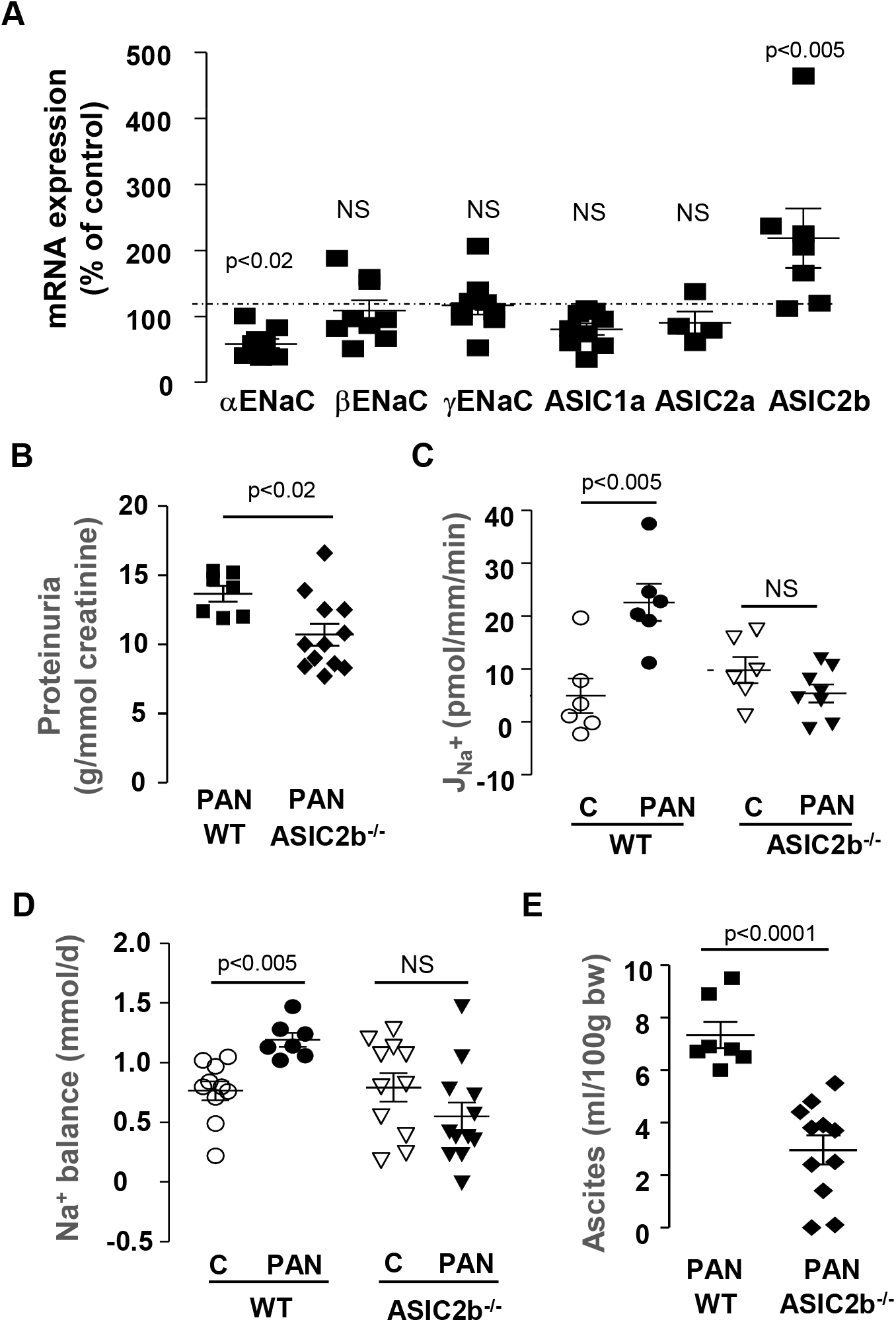
Role of ASIC2b in sodium retention in CC-PAN rats. **A.** RT-qPCR analysis of ENaC/degenerin mRNA in CCDs from CC-PAN rats. Data are expressed as percent of values in corticosteroid-clamped control (CC-Control) rats. Each value represents a rat. **B.** Proteinura in wild type (WT) and ASIC2b^−/−^ CC-PAN rats. Proteinuria is expressed as a function of creatinine excretion. Each value represents a rat. **C.** J_Na_+ in CCDs from WT and ASIC2b^−/−^ rats under basal (C) or nephrotic (PAN) conditions. Each value represents a rat. **D.** Urinary sodium balance in WT and ASIC2b^−/−^ rats under basal (C) or nephrotic (PAN) conditions. Each value represents a rat. **E.** Volume of ascites, as a function of body weight, in WT and ASIC2b^−/−^ CC-PAN rats. Each value represents a rat.

### Molecular characterization of a new variant of ASIC2b in CC-PAN rats

ASICs are mainly expressed in the nervous system where their activation by extracellular acidification induces very brief cation currents that depolarize the cell membrane, allowing activation of nearby voltage-dependent channels or release of neuro-mediators (Deval *et al*, 2010; Kellenberger & Schild, 2015; Lingueglia, 2007). Given the transiency of ASIC-driven currents, ASICs cannot sustain epithelial sodium reabsorption. Therefore, we searched for a variant of ASIC2b that could convert ASIC1a or ASIC2a into a channel carrying sustained sodium current. Starting from RNAs extracted from a CC-PAN rat kidney, we generated by 5’-RACE a cDNA the sequence of which was 100% identical to the deposited rat ASIC2b sequence (ACCN1, variant 1; NM_012892) except that it lacked the 207 5’-most nucleotides (figure 3A). Interestingly, this truncated sequence (GenBank deposition # KP294334) contains an ATG triplet (starting in position 22) within the same reading frame as the translation initiating codon of the full length sequence, suggesting that this short sequence might be translated into a truncated variant of ASIC2b (t-ASIC2b) lacking the first 71 N-terminal amino acid residues, i.e. most of the N-terminal intracellular domain of ASIC2b (figure 3B). Heterologous expression of the corresponding short cRNA in either OKP or HEK cells demonstrated that it is indeed translated into a protein of reduced molecular weight (figure 3C).

**Figure 3.**
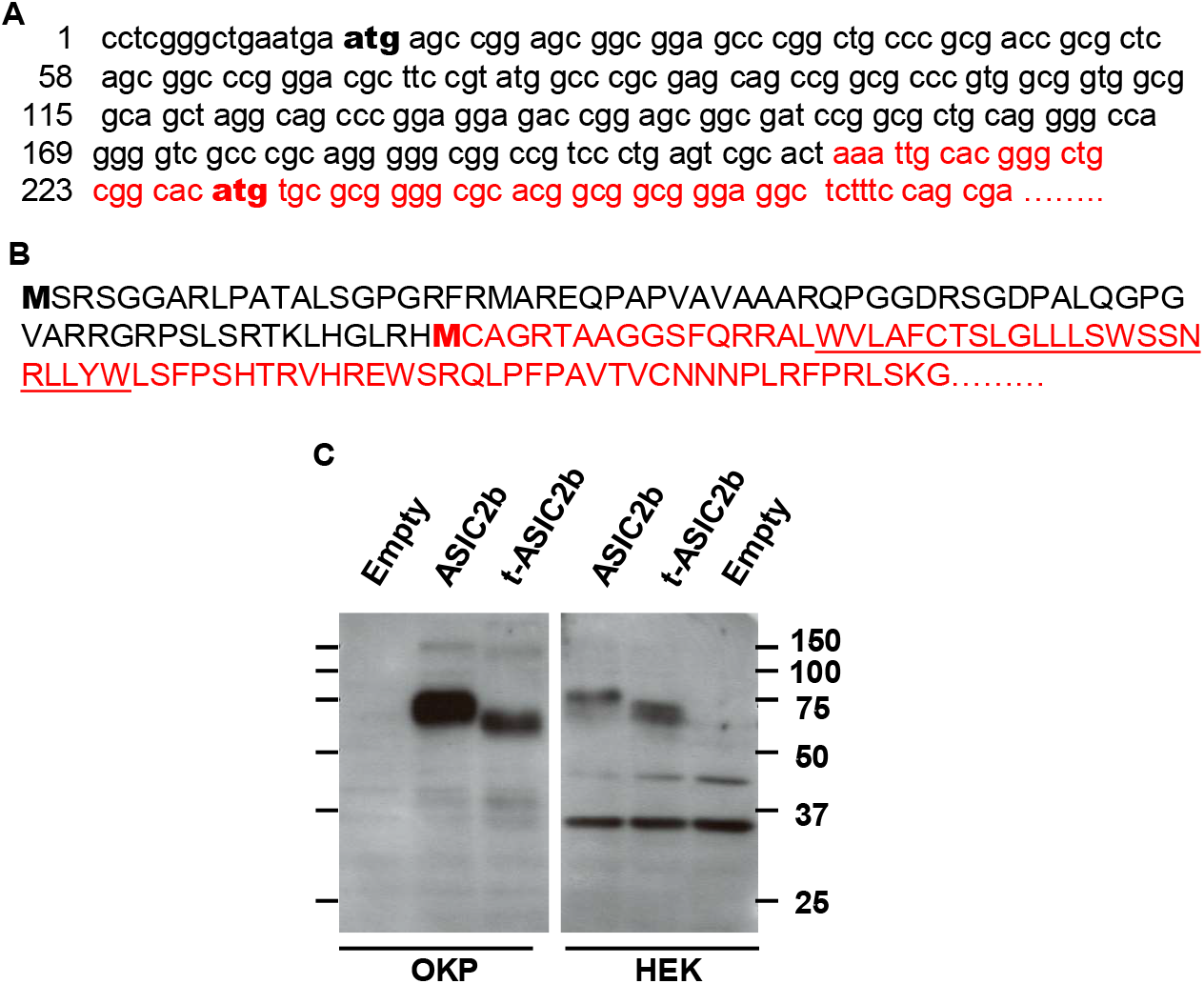
Long and truncated variants of ASIC2b. **A**. 5’ terminal sequence of rat ASIC2b cDNA (Accn1, transcript variant MDEG2, NM_012892, black) and of the short cDNA cloned from CC-PAN CCD (red, GenBank # KP294334). The short sequence contains a putative translation initiation codon in frame with the ASIC2 ATG (in bold). **B**. N-ter sequence of rat ASIC2b (black) and its putative truncated variant (red). The underlined sequence shows the first transmembrane domain. **C.** Western blot analysis of ASIC2 expression in OKP and HEK cells transiently transfected with ASIC2b or its truncated variant (t-ASIC2b) or an empty vector.

We next looked for expression of t-ASIC2b and of ASIC1a and ASIC2a in the CCD of CC-PAN rats. Immunobloting with a specific anti-ASIC1a antibody (Sigma, anti-ACCN2, sab2104215) failed to reveal the expression of ASIC1a in the kidney of CC-PAN rats (not shown). ASIC2a and ASIC2b are produced by alternative splicing of the same ACCN1 gene; they differ by their N-terminal domain consisting of 185 and 236 isoform-specific amino acid residues for rat ASCI2a and ASIC2b respectively. An anti-ASIC2a specific antibody (Alomone, Anti-ASIC2a, ASC-012) revealed a single band ~80kDa in kidney from CC-PAN rats. The same band was found in CCDs and its intensity was increased in CC-PAN rats as compared with CC-controls (Figure 4A), suggesting an over expression of ASIC2a although its mRNA level was unchanged. An ASIC2b-specific antibody directed against an epitope present in full length ASIC2b but not in t-ASIC2b (Interchim, Anti-rat MDEG2, MDEG21-a) revealed no immunoreactivity in CC-PAN rat kidneys (not shown) suggesting that the full length ASIC2b protein is not expressed in the kidney of CC-PAN rats. In absence of commercially available antibody specific for the t-ASIC2b protein, we used a pan anti-ASIC2 antibody directed against an epitope common to ASIC2a and ASIC2b (Abcam, Anti-ACCN1, ab77384). This antibody revealed a faint band ~80kDa (where ASIC2a was detected with the specific antibody) and a main band ~55kDa which was absent in the kidney of ASIC2b^−/−^ rats indicating the presence of a t-ASIC2b protein in CC-PAN rat kidneys. The intensity of this band was higher in CCDs from CC-PAN rats as compared with CC-controls (Figure 4B).

**Figure 4.**
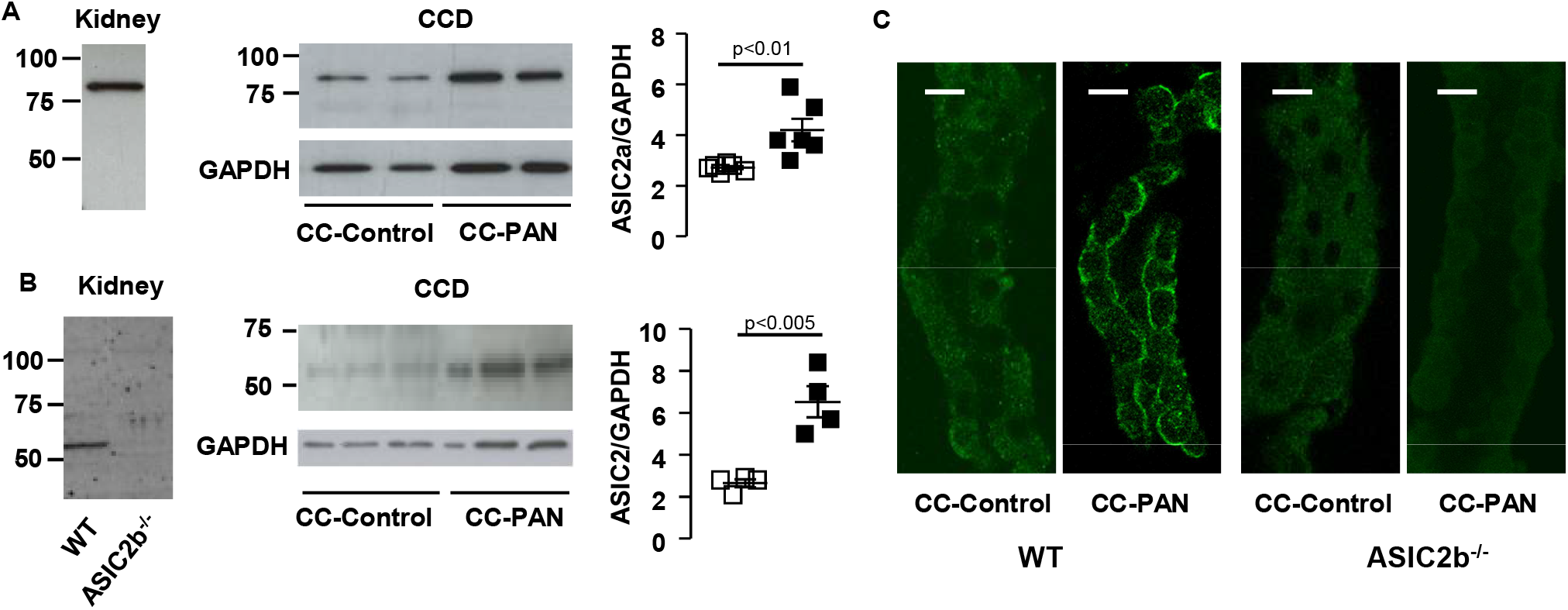
Renal expression of ASIC2. **A and B.** Western blot analysis of ASIC2 expression in kidney of WT and ASIC2b^−/−^ CC-PAN rats and in CCDs of CC-Control and CC-PAN rats using a specific ASIC2a antibody (A) or a pan-ASIC2 antibody (B). Left, representative blot; right, densitometric analysis. Each value represents a rat. **D.** Immuno-labeling of isolated CCD from WT and ASIC2b^−/−^ CC-Control and CC-PAN rats with a pan-ASIC2 antibody. Bar: 10μm

Immuno-histological labeling of isolated CCD with the pan-ASIC2 antibody demonstrated luminal expression of ASIC2a/b in CC-PAN rats but neither in CC-control rats nor in CC-control or CC-PAN ASIC2b^−/−^ rats (figure 4C). Conversely to ENaC which is specifically expressed in principal cells, ASIC2 labeling was detected in all cells, i.e. principal and intercalated cells.

### Functional characterization of t-ASIC2b/ASIC2a channels in X. laevis oocyte

We next evaluated whether t-ASIC2b is functional and may alter ASIC2a properties by expression in *X. laevis* oocytes. Two-electrode voltage clamp experiments showed that expressing t-ASIC2b alone did not induce any acid-sensitive ion current in oocytes (not shown) whereas its co-expression with ASIC2a modified the acid-induced current mediated by the latter. Figure 5A shows representative traces obtained at a holding potential of −70mV in oocytes expressing ASIC2a alone or combinations of ASIC2a with either ASIC2b or t-ASIC2b. As previously described (Ugawa *et al*, 2003), decreasing the extracellular pH from 7.4 to 4.0 rapidly induced a transient current that spontaneously inactivated almost completely within seconds in ASIC2a-expressing oocytes. Co-expression of ASIC2a with the full length ASIC2b increased the initial (peak) current and decreased the level of inactivation, leading to an enlarged residual (plateau) current. These changes were markedly amplified by the co-expression of t-ASIC2b with ASIC2a. The ratio of the plateau over the peak currents was significantly higher in oocytes co-expressing ASIC2a and t-ASIC2b as compared with those expressing ASIC2a alone or in combination with ASIC2b (Figure 5B). In oocytes co-expressing ASIC2a and t-ASIC2b, the residual desensitized current remained stable for >5 min (not shown). These findings indicate that co-expression of ASIC2a with a N-ter truncated form of ASIC2b induces the formation of functional channels that allow sustained transport of sodium in response to an acid stimulus. In the following experiments, we focused our analysis on the residual current carried by the desensitized channel, the one which can account for sodium reabsorption in CCDs.

**Figure 5.**
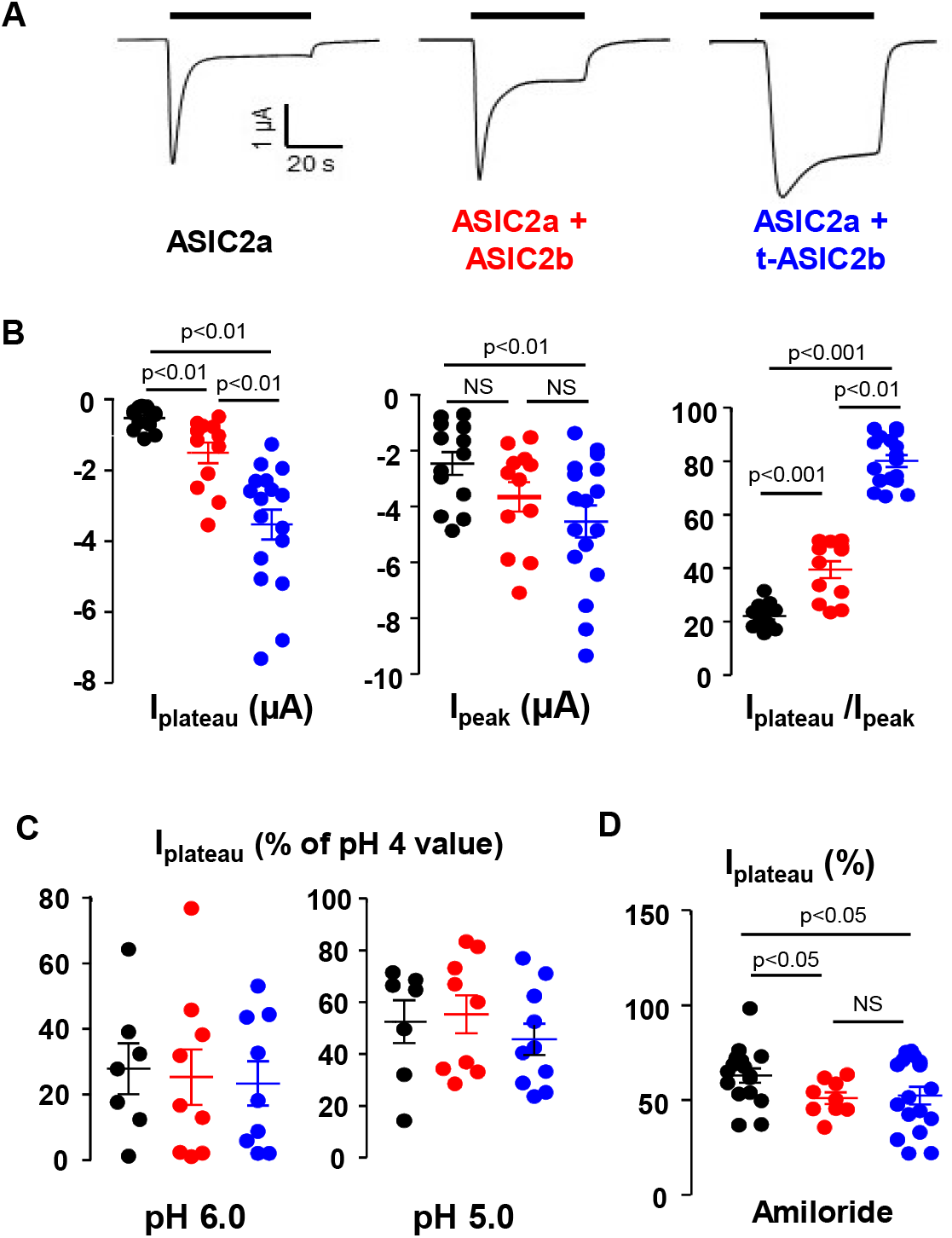
Functional expression of ASIC2 in *X. laevis* oocyte. **A.** Asic2b co-expression attenuates Asic2a desensitization. Original traces of whole cell current (holding potential = −70 mV) in oocytes expressing ASIC2a alone or with ASIC2b, or t-ASIC2b, as indicated below the traces. Inward currents were induced by a rapid extracellular acidification from pH 7.4 to pH 4.0 (indicated by the horizontal bar). **B.** Mean transient (peak) and residual (plateau) induced-currents (corrected by currents recorded in control oocytes) in oocytes expressing ASIC2a alone (black bars) or co-expressing ASIC2b (red) or t-ASIC2b (blue). Each point represents an oocyte. **C.** Acid-induced plateau currents (normalized to maximal value achieved at pH 4) at different extracellular pH; colors as above. Each point represents an oocyte. **D.** Effect of amiloride: Acid-induced (pH 4) plateau currents (normalized to the value measured in the absence of inhibitor) in the presence of 100 μM amiloride. Each point represents an oocyte.

Within the range of pH tested (7.4 to 4.0), the pH sensitivity of the plateau current was similar in oocytes injected with ASIC2a alone or in combination with ASIC2b or t-ASIC2b (Figure 5C). Truncated ASIC2b did not alter significantly the apparent low sensitivity of ASIC2a to amiloride (Figure 5D). Co-expression of t-ASIC2b changed neither the sodium affinity nor the cation selectivity of ASIC2a (not shown).

### Expression of ASIC2a/b in nephrotic patients

We next evaluated whether ASIC2a/b is also expressed in the kidney of patients with minimal change disease. RT-qPCR revealed the presence of ASIC2a/b mRNA in kidney biopsies from 5/8 patients with idiopathic nephrotic syndrome whereas it was undetectable in the remaining 3 (figure 6A). All patients with other renal diseases (26/26) displayed no or very low expression of ASIC2a/b mRNA. Immunohistochemistry showed ASIC2a/b labeling in collecting ducts of some, but not all, nephrotic patients whereas labeling was never observed in non-nephrotic patients (figure 6B).

**Figure 6.**
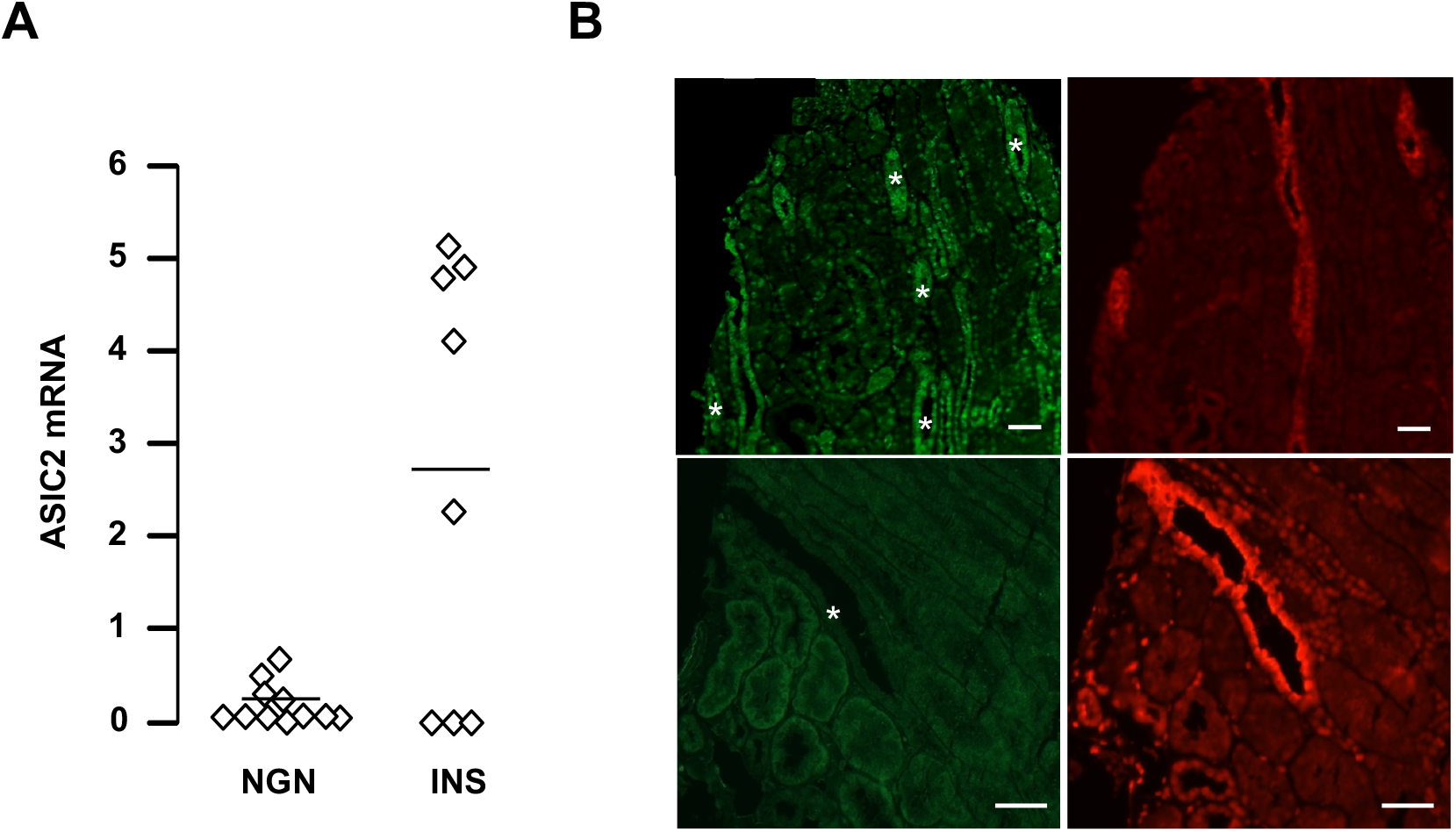
Expression of ASIC2 in nephrotic patients. **A.** RT-qPCR analysis of ASIC2 mRNA in kidney biopsies from patients with non glomerular nephropathies (NGN, n=11) or idiopathic nephrotic syndrome (INS, n=8). Given the heterogeneity of biopsies in term of cell composition, data were standardized using distal nephron markers, as previously described (Disset *et al.*, 2009). Data are in arbitrary units. **B.** Immunolabeling of kidney serial sections with anti-AQP2 (a & c, red) and anti-ASIC2 antibody (b & d, green) from two patients with idiopathic nephrotic syndrome (a, b) or a non-nephrotic patient (c, d). Bar: 100μm.

### Signaling of ASIC2b induction during nephrotic syndrome

It has been reported that albumin abnormally present in the kidney ultrafiltrate of nephrotic rat is endocytosed in CCDs, and that this process triggers several intracellular signaling cascades (Dizin *et al*, 2013; Fila *et al*, 2011). We therefore investigated the role of albumin in the induction of ASIC2b in CC-PAN rats. For this purpose, we used Nagase analbuminemic rats (NAR), a strain spontaneously lacking the albumin gene. Despite analbuminemia, CC-NARs developed massive but slightly lower proteinuria than control rats in response to PAN (Figure 7A). This proteinuria mainly consisted of proteins of higher molecular weight than albumin (Figure 7B). In CC-NARs, PAN increased neither ASIC2b mRNA expression in CCD nor sodium retention (Figure 7C & 7D). The volume of ascites was reduced by half in NARs as compared to WT rats (in ml/100g body wt ± SE; WT: 7.3 ± 0.5, n=6; NAR: 3.8 ± 0.5, n=6: p<0.001). Interestingly, J_Na_+ in CCDs from nephrotic NARs was not increased compared with non-nephrotic NARs (Figure 7E), indicating that albumin participates in the stimulation of ASIC2-dependent Na^+^ reabsorption.

**Figure 7.**
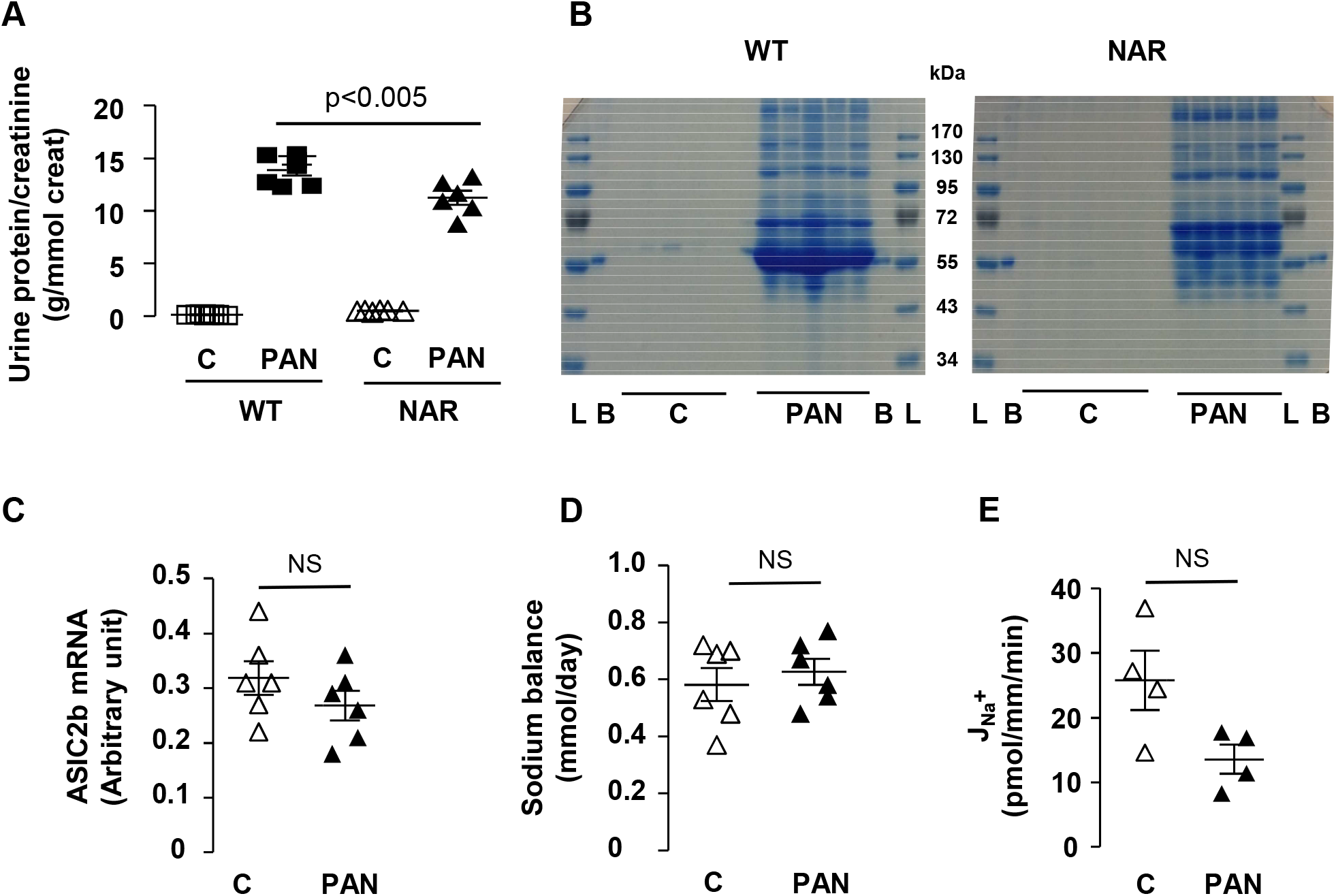
Role of albumin in ASIC2b expression and sodium retention. **A.** PAN-induced proteinuria in CC-WT and Nagase analbuminemic rats (NAR). Proteinuria is expressed as a function of creatinine excretion. Each point represents a rat. **B.** Urinary proteinogram in WT and NAR rats under control (C) and nephrotic conditions (PAN). L, molecular weight markers; B, bovine serum albumin. **C.** Expression of ASIC2b mRNA in control and PAN nephrotic CC-NARs. Each point represents a rat. **D.** Sodium balance in control and PAN nephrotic CC-NARs Each point represents a rat. **E**. J_Na_+ in CCDs from control and PAN nephrotic CC-NARs. Each point represents a rat.

We next evaluated the possible involvement of ERK pathway in the induction of ASIC2b because we previously reported that this pathway is activated by albuminuria (Fila *et al.*, 2011). By semi-quantitative immunofluorescence, we confirmed that phospho-ERK labelling was increased in CCDs from CC-PAN rats as compared with CC-rats, and that this effect was abolished in NARs (Figure 8A), indicating that ERK phosphorylation is induced by albumin, probably through its endocytosis. ERK phosphorylation was also prevented by *in vivo* treatment of CC-PAN rats with the mitogen-activated protein kinase kinase inhibitor U0126 (Figure 8B). U0126 treatment also abolished the induction of ASIC2b mRNA expression and the positivation of sodium balance (Figure 8C-D), and reduced by over half the volume of ascites (in ml/100g body wt ± SE; Control: 7.3 ± 0.5, n=6; U0126: 3.0 ± 0.4, n=5: p<0.001). We also found that U0126 prevented the over-expression of the two subunits of Na,K-ATPase, the motor for sodium reabsorption in CCD (Figure 8D). Unfortunately, we were not able to dissect native CCDs from U0126-treated rats to measure J_Na_+ by *in vitro* microperfusion in these rats.

**Figure 8.**
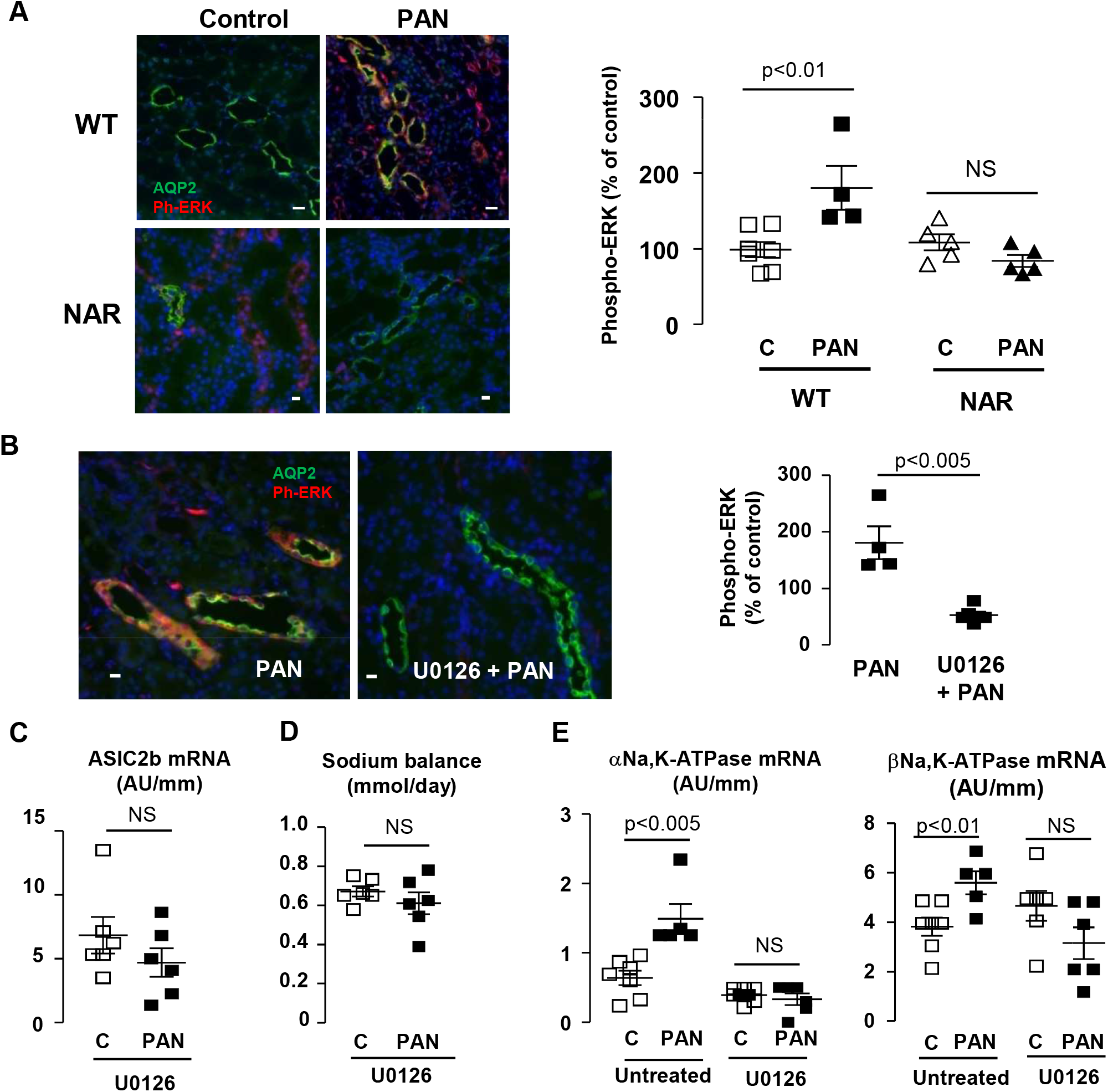
Role of ERK pathway in ASIC2b expression and sodium retention. **A.** Immuno-labeling of kidney cortical sections from CC-WT rats and CC-NARs under control and PAN nephrotic with anti-phospho-ERK (red) and anti-aquaporin 2 (AQP2, in green). AQP2 was used as a marker of CCDs. Left panel: representative images showing phosphorylation of ERK in nephrotic WT rats but not in NARs. Right panel: quantification of phosphor-ERK labeling in CCDs. Values were normalized to the labeling measured in each experiment in the same CC-control WT rat. Each point represents a rat. **B.** Same immune-labeling as in A in CC-PAN WT rats treated or not with the ERK kinase inhibitor U0126. **C.** Expression of ASIC2b mRNA in U016-treated rats under control and nephrotic conditions. Each point represents a rat. **D.** Sodium balance in in U016-treated rats under control and nephrotic conditions. Each point represents a rat. **E.** Expression of mRNAs of the α and β subunits of Na,K-ATPase in U016-untreated and -treated rats under control and nephrotic conditions. Each point represents a rat

Altogether, these results indicate that nephrotic albuminuria activates the ERK pathway and subsequently induces the expression of ASIC2b and sodium retention in CCDs. This mechanism is specific for albumin. This conclusion raises a question regarding the reversal of sodium retention in PAN rats. As a matter of fact, it has been shown than within 12 days following the administration of PAN, sodium balance and Na,K-ATPase activity in CCD return to basal levels and that ascites disappears despite the maintenance of massive proteinuria. We confirmed that within 12 days sodium balance was restored to basal level and ascites was reduced while proteinuria remained high (Figure 9A). We observed that at that time albumin was no longer accumulated in CCDs (Figure 9B) and that ERK phosphorylation and ASIC2b mRNAs had turned back to basal level (Figure 9C-D).

**Figure 9.**
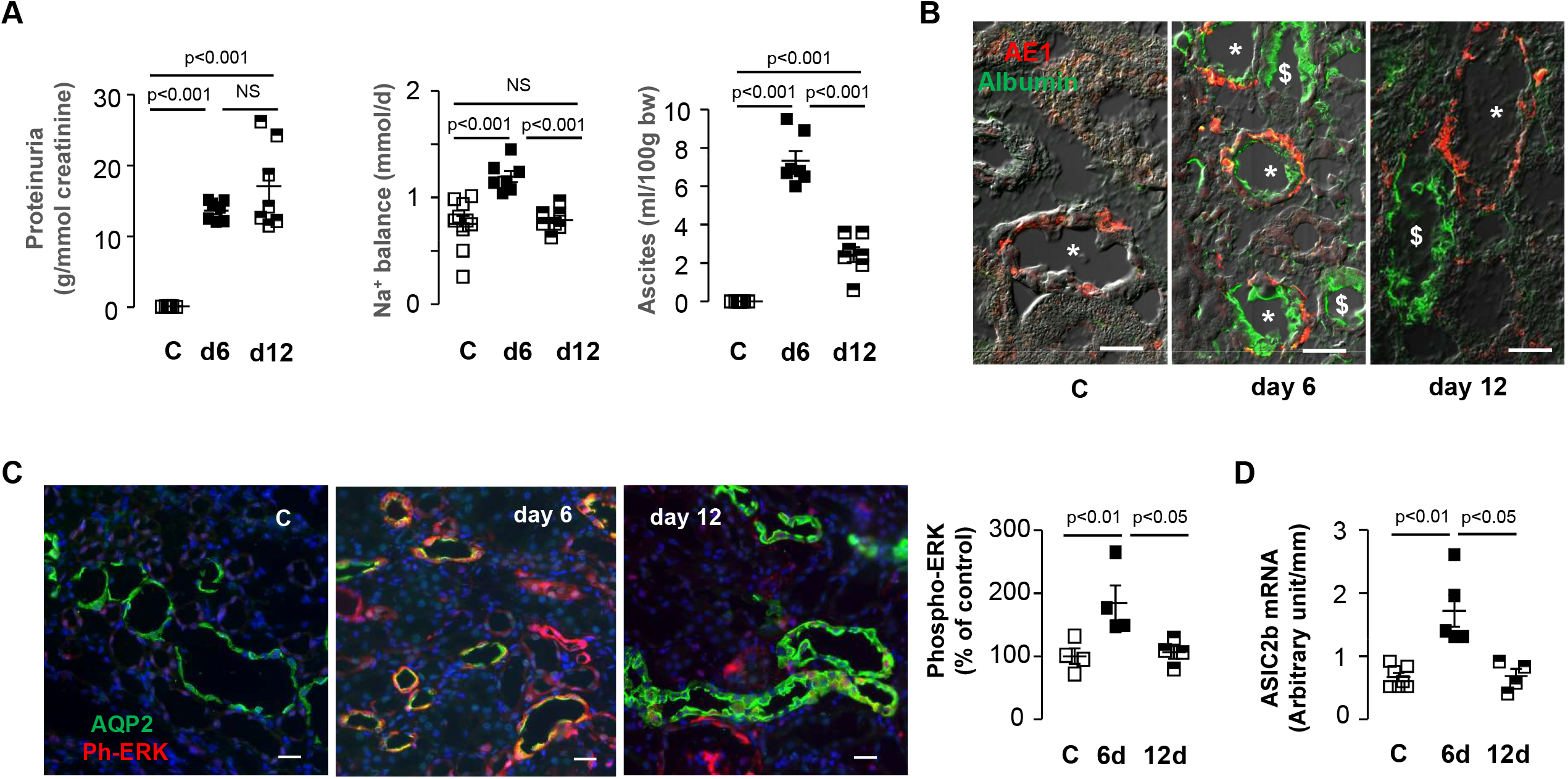
Reversal of sodium retention. **A.** Proteinuria, sodium balance and volume of ascites in CC-rats under control condition (C) or 6 or 12 days after PAN administration. Each point represents a rat. **B**. Immuno-labeling of kidney cortex section from CC-rats under control condition (C) or 6 or 12 days after PAN administration with an anti-albumin antibody (green) and anti-anion exchanger 1 (AE1) antibody (red), a specific marker of CCD intercalated cells. *, CCD; ^$^ proximal tubule. Bars: 25μm. **C.** B.Immuno-labeling of kidney cortex section from CC-rats under control condition (C) or 6 or 12 days after PAN administration with an anti-phospho ERK antibody (red) and an anti-AQP2 antibody (green), a specific marker of CCD principal cells. Left, representative images; right, quantification as in figure 8A. Each point represents a rat. **D.** Expression of ASIC2b mRNA in CCDs from CC-rats under control condition (C) or 6 or 12 days after PAN administration. Each point represents a rat.

## Discussion

Sodium reabsorption in the CCD proceeds via two pathways: the classical electrogenic pathway mediated by amiloride-sensitive ENaC on the apical side and basolateral Na,K-ATPase, and an electroneutral, thiazide-sensitive pathway energized by the basolateral H-ATPase and requiring the concerted activity of apical sodium dependent- and -independent chloride/bicarbonate exchangers and of a basolateral sodium-bicarbonate cotransporter (Chambrey *et al*, 2013; Leviel *et al.*, 2010). These two pathways originate from principal cells and B-type intercalated cells respectively. In contrast, A-type intercalated can secrete sodium via basolateral Na/K/Cl co-transporter and apical H(Na),K-ATPase (Edwards & Crambert, 2017; Morla *et al*, 2016). Here we show that sodium reabsorption in CCD from CC-PAN rats is electrogenic and amiloride-sensitive but independent of any ENaC subunit, since none of them is expressed at the apical cell border (Lourdel *et al.*, 2005), revealing the existence of a third pathway for sodium reabsorption. This pathway involves a newly characterized truncated variant of ASIC2b which, in association with ASIC2a, can carry sustained apical sodium entry. This channel is expressed at the apical border of all cell types constituting CCDs suggesting that they might all participate in sodium reabsorption. However, this appears unlikely for A-type intercalated cells because they do not express any known sodium pump at their basolateral side.

Based on co-immunoprecipitation and functional studies, Ugawa and coworkers have shown that rat ASIC2a and ASIC2b biochemically assemble to constitute functional (Ugawa *et al.*, 2003) channels. The variant of ASIC2b identified in this study differs from ASIC2b by the deletion of most of its intracellular N-terminal domain (71/88 amino acid residues). The N-terminal domain of ASICs is not necessary for subunit association since the trimeric structure of chicken ASIC1 was deduced from crystal structures obtained from proteins with truncated intracellular domains (Dawson *et al*, 2012; Jasti *et al*, 2007). This indicates that ASIC2a and truncated ASIC2b can assemble to constitute functional channels likely made of three subunits like all functional ASICs. However, the stoichiometry of assembly remains unknown and may be variable *in vivo* as well as in our co-expression studies. Consequently, the macroscopic currents measured in *X. laevis* oocyes may stem from a mix of three types of trimeric channels containing 0, 1 or 2 subunits of t-ASIC2b.

Current kinetics models propose that ASICs exist under three functional states: a closed, an open and a desensitized state in which the channel is partially open but cannot be activated by acid stimulus. Following acid stimulus, the open and desensitized states translate into peak and plateau currents respectively. It was previously reported that the main effect of ASIC2b is to increase the current carried by desensitized ASIC2a (Ugawa *et al.*, 2003). We confirmed this effect of ASIC2b and found that it is amplified by t-ASIC2b. Thus, the kinetics of the inward current mediated by ASIC2a/t-ASIC2b resembles that of ENaC-mediated current, except for the presence of a transient peak of small amplitude as regards to the remaining plateau. Another important issue concerns the intensity of the current carried by ASIC2a/t-ASIC2b compared with ENaC-mediated current. Formally, this question cannot be answered by electrophysiological analysis in *X. laevis* oocyte since the current intensity depends on the expression level of the respective channels and possibly on the stoichiometry of t-ASIC2b/ASIC2a assembly. Nonetheless, it is worth mentioning that the macroscopic Na^+^ current measured in ASIC2a and t-ASIC2b-expressing oocytes under the desensitized state (~3μA at a holding potential of −70mV) is of the same order of magnitude as that initially reported in ENaC-expressing oocytes (~1μA at −100mV) (Canessa *et al*, 1994). Altogether our findings indicate that, given their conductive properties under desensitized state, hetero-trimers made of ASIC2a and t-ASIC2b may substitute for ENaC and allow sustained sodium reabsorption in CCD principal cells.

Conversely to ENaC which opens stochastically in absence of stimulus, ASIC2 requires an acid stimulus to open and to desensitize. Thus, the increased abundance of ASIC2 subunits in CCDs from CC-PAN rats is not sufficient to account for sodium reabsorption; this also requires the presence of an ASIC2 activating factor. ASIC2a requires low pH for maximal activation (pH_50_ ~4.0, maximal current, pH~2.0) (Ugawa *et al.*, 2003), and association with t-ASIC2b did not modify this pH dependency. The pH of the luminal fluid prevailing in CCDs *in vivo* is estimated in the 6.0-6.5 range (DuBose *et al*, 1979; Weinstein, 2010) but, because type A intercalated cells of the CCD secrete protons and due to the presence of unstirred layers, the pH at the external surface of the apical membrane might be 0.5-1.0 units lower, i.e. in the 5.0-6.0 range. Based on *X. laevis* expression experiments, this pH would induce ~10% of the current carried by desensitized ASIC2a/t-ASIC2b at pH 4.0, which may be sufficient to account for the rate of sodium reabsorption determined by *in vitro* microperfusion. In addition, one cannot exclude that factors abnormally present in the urine of nephrotic rats or produced by CCD cells may also increase the pH sensitivity of ASIC2a/t-ASIC2b or activate it independently of pH. Such mechanisms have been reported with other ASICs, e.g. NO and arachidonic acid potentiate ASIC-mediated proton-gated currents (Cadiou *et al*, 2007; Smith *et al*, 2007) and the arginine metabolites agmatine and arcaine activate ASIC in a proton-independent manner (Li *et al*, 2010; Yu *et al*, 2010). Moderate proteolysis of ASIC2 may also activate it, as previously reported for ENaC and plasmin (Passero *et al*, 2008), the excretion of which is increased during nephrotic syndrome (Svenningsen *et al*, 2009).

The apparent low amiloride-sensitivity of t-ASIC2b/ASIC2a observed in *X. laevis* oocytes (IC_50_≈50 μM) contrasts with the high sensitivity of sodium transport observed *in vitro* in microperfused CCDs (full inhibition with 10 μM amiloride). These differences in amiloride sensitivity likely stem from the fact that acid pH required to activate ASIC2 in *X. laevis* oocytes strongly decreases its sensitivity to amiloride (Ugawa *et al.*, 2003). This further support the notion that acid pH is probably not the unique activating factor of ASIC2 in ASDN during nephrotic syndrome.

The mechanism of sodium retention in nephrotic rats varies according to their aldosterone status: when animals display high plasma aldosterone level, increased sodium reabsorption is mediated by the classical ENaC-dependent pathway and there is no evidence for ASIC2 expression, whereas blunting of hyper aldosteronemia switches off the ENaC pathway and triggers the ASIC2-dependent one. This suggests the existence of a balance between factors that reciprocally trigger and repress the renal expression of ENaC and ASIC2. Aldosterone is the major inducer of ENaC expression in the ASDN; whether or not it represses the expression of ASIC2 remains to be established. Here we report that the phosphorylation of ERK brought about by the endocytosis of albumin mediates the over expression of ASIC2 in the ASDN and the stimulation of sodium transport. In contrast, ERK phosphorylation reduces ENaC activity by different mechanisms including the decrease of its open probability, increasing its membrane retrieval (Falin *et al*, 2005) and decreasing expression of its mRNA expression (Niisato *et al*, 2012). Thus, endocytosis of albumin and subsequent activation of ERK pathway appears as a major factor that switches on and off ASIC2 and ENaC pathways respectively. This raises the question of the mechanism responsible for the escape of sodium retention to albuminuria observed in the long term. Our data show that it proceeds at the step of albumin endocytosis, possibly via the down regulation of the albumin receptor 24p3R (Dizin *et al.*, 2013).

Our preliminary results in nephrotic patients indicate that expression of ASIC2 in the ASDN is not restricted to the PAN rat model. The fact that only part of the patients studied displayed ASIC2 expression in their ASDN might be related to the fact that, in nephrotic rats, ASIC2 is only found in corticosteroid clamped animals and that only a fraction of nephrotic patients displays normal aldosterone status (Vande Walle *et al.*, 1995). A further study will be necessary to evaluate whether expression of ASIC2 in nephrotic patients is correlated with low plasma aldosterone level. More generally, it is noteworthy that the alternate ATG initiating codon found in the rat ASIC2b sequence is conserved not only in the human sequence (NM_183377.1) but also in mice (NM_007384.3).

It has been reported previously that channels made of αENaC and ASIC1a constitute functional cation reabsorbing channels that participate in fluid clearance by lungs (Trac *et al*, 2017). Here we describe for the first time the functional expression and role of channels exclusively made of ASIC subunits out of excitable cells. It also shows that deletion of the intracellular N-terminal domain of ASIC2b modifies its properties and allows converting the transient ASIC2a into a long lasting epithelial channel. Thus, ASICs and ENaC share more than sequence similarities since both can perform epithelial sodium transport. Whether variants of ENaC might work as transient sodium channels is a stimulating hypothesis.

## Methods

### Animals

The animals were kept at CEF (Centre d’Explorations Fonctionnelles of the Cordeliers Research Center, Agreement no. B75-06-12). All experimental protocols were performed in accordance with the institutional guidelines and the recommendations for the care and use of laboratory animals put forward by the Directive 2010/63/EU revising Directive 86/609/EEC on the protection of animals used for scientific purposes which are equivalent of the ARRIVE guidelines and were approved by the Charles Darwin Ethic Committee (#Ce5-2012-041). Experiments were carried out on male rats (150–170 g at the onset of the experimentation) fed a standard laboratory chow (A04, Safe, Augy, France) with free access to water. Sprague–Dawley rats were from Charles Rivers (L’Abresles, France) and Nagase analbuminemic rats (NARs) were from Japan SLC (Shizuoka, Japan). For surgery, animals were anaesthetized by intraperitoneal injection of a mix including Domitor (Pfizer, 0.5 *μ*g/g body wt), Climasol (Graeub, 2 *μ*g/g body wt) and Fentanyl Jansen (Janssen Cilag Lab, 5 ng /g body wt). Animals were awakened by a subcutaneous injection of a mix containing Antisedan (Pfizer, 750 ng/g body wt), Sarmasol (Graeub, 200 ng/g body wt) and Narcan (Aguettant, 133 ng/g body wt). Before killing, animals were anaesthetized with pentobarbital (Sanofi, France, 50 mg/kg body wt, I.P.). Corticosteroid clamp was achieved by bilateral adrenalectomy and supplementation with aldosterone (10 μg/kg/day) and dexamethasone (14 μg/kg/day) through subcutaneous osmotic pump (ALZET, Charles River) (Lourdel *et al.*, 2005) Nephrotic syndrome was induced the day after surgery for corticosteroid clamp by a single intra-jugular injection of aminonucleoside puromycine (PAN) (Sigma-Aldrich, 150 mg/kg body wt). Control rats received a single injection of isotonic NaCl (1ml /100 g body wt). A group of rats was treated with the ERK kinase inhibitor U0126 by daily subcutaneous injection (3mg/10 g body wt/day in a mixture of DMSO/sesame oil, 16%/84% vol/vol). Treatment started the day of corticosteroid clamp. Animals were studied 6 days after vehicle or PAN injection, at the time of maximum of sodium retention and proteinuria, or after 12 days when sodium balance was restored (Deschenes & Doucet, 2000). To induce ENaC expression, rats were fed a Na^+^-depleted diet (Safe, synthetic diet containing 0.11g Na^+^/kg instead of 2.5g/kg). Rats were studied 14 days after the onset of the low Na^+^ diet.

#### Generation of ASIC2b null rats

Invalidation of *Accn1* gene encoding the ASIC2 proteins was performed using Crispr-Cas9 technology (TRIP, INSERM UMR1064, Nantes, France). Zygotes from Sprague-Dawley rats were microinjected with a single guide RNA (sgRNA, 10ng/ μl) designed to target exon 1 of the *Accn1* gene and Cas9 mRNA (50ng/μl) as previously described (Menoret *et al*, 2015).This resulted in a deletion of 20 nt and a frameshift of the coding region leading to the early appearance of a premature stop codon. Embryos were then implanted in pseudopregnant females and grown until full term. To genotype the animals, a 953 pb genomic DNA fragment around the targeted region was PCR to detect the deleted fragment in KO animals by gel electrophoresis.

### Human material

Serial sections of frozen human kidney biopsies for RNA extraction or fixed human kidney biopsy for immunohistochemistry were provided by the pathology department at Robert Debré Hospital. Inform consent of each patient was collected according to the French current regulation. cDNA samples from kidney biopsies of non-nephrotic patients were from a previously published series of experiments (Disset *et al*, 2009).

### Urine proteinogram

Urine samples were thawed and centrifuged 10 min at 3300g. 9 μl of supernatant (pure for control rats, 1/10 diluted in H_2_O for PAN rats) were half diluted in Glycerol/blue solution and separated by SDS-PAGE. After migration gels were rinsed twice with water and stained overnight in Coomassie blue solution (Pageblue protein staining, Lifetechnologies) at room temperature. The gels were then rinsed for at least 48h in water.

### Microdissection of cortical collecting ducts (CCDs)

CCDs were dissected under stereomicroscopic observation either from fresh kidney slices (microperfusion) or after liberase treatment (Blendzyme 2, Roche Diagnostics, Meylan, France) (immunoblotting, RT-qPCR and immuno-histochemistry) (Fila *et al.*, 2011). For RT-qPCR experiments, microdissection was performed under RNase-free conditions.

### In vitro microperfusion

Unless indicated otherwise, CCDs were perfused at 37 °C under symmetrical conditions (Vande Walle *et al.*, 1995), with bath and perfusate containing (in mM): 138 NaCl, 1.2 MgSO_4_, 2 K_2_HPO_4_, 2 calcium lactate, 1 sodium citrate, 5.5 glucose, 5 alanine, 12 creatinine (bath continuously gassed with 95% O_2_/5% CO_2_). pH was adjusted either to 7.4 with Hepes 10 mM and Tris 5 mM or to 6.0 with MES 5 mM. Transepithelial voltage (PD_te_) was measured via microelectrodes connected through an Ag/AgCl half-cell to an electrometer. Tubules were perfused at low rate (~2 nl/min). Concentrations of Na^+^, and creatinine were determined by HPLC, and ion flux (JNa^+^) was calculated as:

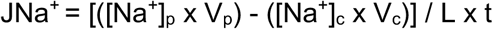

where [Na^+^]_p_ and Na^+^]_c_ are the concentration of Na^+^ in the perfusate and collection, respectively, V_p_ and V_c_ are the perfusion and collection rates, respectively, L is the tubule length, and t is the collection time. For each tubule, Na^+^ flux was calculated as the mean of four successive 10–15min collection periods. V_p_ was calculated as:

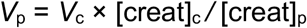

where [creat]_c_ and [creat]_p_ are the concentrations of creatinine in the collection and perfusate, respectively.

### RNA extraction and RT-qPCR

RNAs were extracted from 40–50 CCDs using an RNeasy micro kit (Qiagen, Hilden, Germany) as previously described (Fila *et al.*, 2011). Frozen human kidney biopsies (2-3mg) were dissolved in 5μl of RLT complemented with β-mercaptoethanol, transferred in a tube and RNAs were extracted as above. RNAs were reverse transcribed using first strand cDNA synthesis kit for RT-PCR (Roche Diagnostics, Meylan, France), according to the manufacturers’ protocols. Real time PCR was performed with a LightCycler 480 SYBR Green I master quantitative PCR kit (Roche Diagnostics) according to the manufacturer’s protocol, except that reaction volume was reduced 2.5-fold. Specific primers (table 1) were designed using Probe Design (Roche Diagnostics).

**Table 1.**
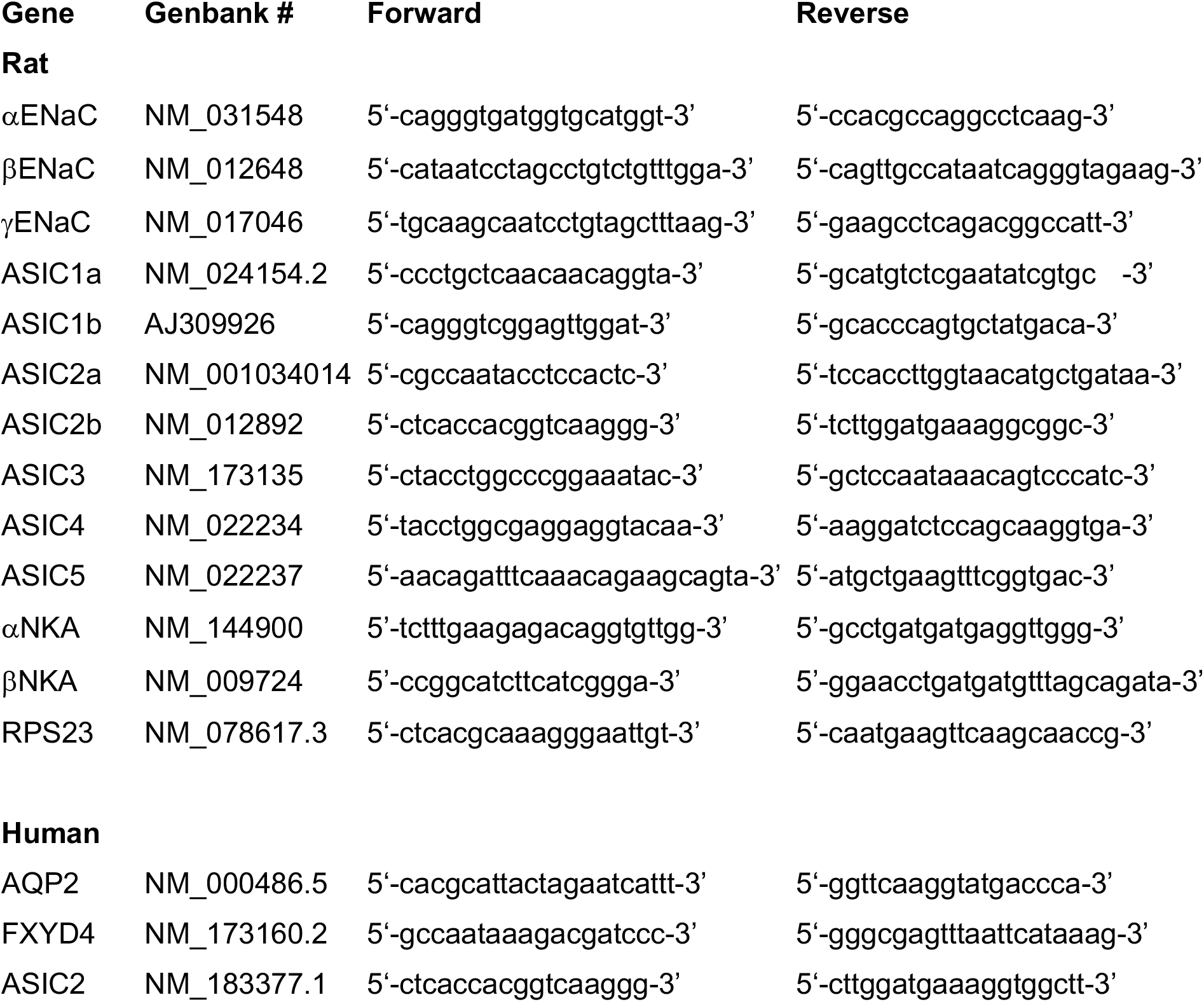
Sequence of primers used for qPCR.

In each experiment, a standardization curve was made using serial dilutions of a standard cDNA stock solution made from rat or human kidney, and data (in arbitrary unit) were calculated as function of the standard curve. For CCDs, results were normalized as a function of Rps23 expression. Data are means ± SE from several animals. Given the heterogeneity of human kidney biopsies in term of cell composition, data were normalized using the distal nephron markers aquaporin 2 (AQP2) and FXYD4, as previously described (Disset *et al.*, 2009).

### 5’-RACE

Total RNA was extracted from the kidney of a corticosteroid-clamped PAN rat following the Tri Reagent protocol (Sigma), Dnase I-treated, and purified using RNeasy minikit (Qiagen) according the manufacturer’s instructions. The 5’-RACE cDNA was synthesized by using the GeneRacer Kit (Invitrogen) according the manufacturer’s instructions. The first PCR reaction was performed using ASIC2b-specific reverse primer (5’- cacaaggagtgtgcagagcctgc-3’) and the GeneRacer 5’ primer supplied with the RACE cDNA kit. The nested ASIC2b-specific reverse primer (5’-cctttcatccaagagctgggctttg-3’) and the GeneRacer 5’ Nested primer supplied with the kit were used for the second PCR reaction. The nested PCR products were subcloned in the pCR4-TOPO vector (TOPO TA cloning Kit for Sequencing, Invitrogen) according the manufacturer’s instructions and sequenced.

### ASIC2 subcloning

The full length rat ASIC2b (accn1, variant 1, NM_012892) was purchased from ImaGenes (Germany) and cloned into pMK (kanR) using SfiI cloning sites. Asic2b N416A and Asic2b N443A mutants were generated using Quickchange Site-Directed Mutagenesis kit (Agilent Technologies) according to the manufacturer’s instructions. For expression in *X. laevis* oocytes, the fragment of interest was extracted and subcloned in pGH19 expression vector Bam HI/ Xba I cut. For expression in eukaryotic cells the pcDNA™ 3.1/V5-His TOPO® TA Expression Kit (Invitrogen) was used according to the manufacturer’s protocol. The Bam HI and Xba I sites were used to subclone the insert of interest.

The truncated variant of rat ASIC2b was obtained as described above and subcloned in EcoRI site into pTLB expression vector for expression in *X.laevis* oocytes. Subcloning for expression in eukaryotic cells was performed as for full length ASIC2b.

Full length rat ASIC2a (accn1, variant 2, NM_001034014) was kindly provided by Dr. E. Lingueglia in a mammalian vector and cloned in pGH19 *X. laevis* expression vector. All constructs were verified by sequencing.

### Expression in mammalian cells

OKP and HEK293 cells were cultured in DMEM medium, high glucose, glutamax (GIBCO, Invitrogen) or DMEM medium, high glucose, Hepes, supplemented with 100mM pyruvate respectively. Both media were supplemented with 10% fetal bovine serum (Invitrogen), penicillin (100 units/ml), and streptomycin (100 μg/ml). Cultures were incubated at 37°C in a humidified 5% CO_2_-air atmosphere. Cells grown for 24h in 6-well plates were transfected with 1 μg of cDNA construct (Asic2b or t-Asic2b) using a Lipofectamine™ Plus kit (Invitrogen) according to the manufacturer’s protocol. Transiently transfected cells were studied 48 h after transfection. Empty vector was used as control.

Cells were solubilized in buffer containing 50mM Tris-HCl pH7.4, 150mM NaCl, 2mM EDTA, 1% (v/v) triton X-100, 0.1% SDS and Complete™ protease inhibitor cocktail (Roche Diagnostics) and proteins were separated by SDS-PAGE and analyzed by western blot using pan-ASIC2 antibody as described for CCDs.

### Functional expression in Xenopus laevis oocytes

Linearized pGH19-ASIC constructs were transcribed *in vitro* from the promoter using a mCAP mRNA mapping kit (mMESSAGE mMACHINE T7/SP6 kit, Ambion, Austin, TX, USA).

Oocytes were obtained following the partial ovariectomy of female adult *X. laevis* (obtained from Xenopus Express, Vernassal, France). The procedure of anaesthesia and surgery was approved by the Paris-Descartes Ethic Committee for Animal Experiments (#CEEA34.GP.011.12). Stage V-VI oocytes were selected after enzymatic defolliculation (2 h of gentle shaking in a collagenase-containing solution (1A, Sigma) at 16°C). Control oocytes were injected with 50 nl of RNase-free water; ASIC-expressing oocytes were injected with 50nl of RNAse-free water containing 3ng of ASIC2a (oocytes expressing ASIC2a alone) or a mixture of 3 ng of ASIC2a cRNA plus 6 ng of ASIC2b (oocytes co-expressing ASIC2a and ASIC2b). Oocytes were incubated for 2-4 days at 16°C before electrophysiological experiments or analysis of protein expression.

Two-electrode voltage clamp was achieved using low resistance (<1 MΩ) glass microelectrodes, filled with 3M KCl, and connected to a voltage-clamp amplifier (Turbo TEC-10CX NPI Electronic, Tamm, Germany). Whole cell currents were recorded at the holding potential VH= −70 mV interfaced to a computer with Digidata 1322A analyzed using Axon pClamp 9 software (Axon Inst, USA). The oocyte was continuously superfused during the experiment with ND 96 solution (in mM: NaCl 96, KCl 2, CaCl_2_ 1.8, MgCl_2_ 1, Hepes 5, pH 7.4) buffered with 5 mM HEPES and adjusted to pH 7.4 or with 5 mM MES and adjusted to pH 4-6 to elicit acid-induced currents.

### Immunoblotting

Pools of 30-80 CCDs were solubilized in Laemmli buffer, and proteins were separated by SDS-PAGE and transferred to polyvinylidene difluoride membranes (GE Healthcare). After blocking, blots were successively incubated with specific anti-ASIC2a antibody (Alomone labs, ASC-012, 1/1000) or pan-ASIC2 antibody (Abcam, ab77384, 1/700) and anti-rabbit IgG antibody coupled to horseradish peroxidase (Promega France, W401B, 1/2500) and revealed with the Western Lightning chemiluminescence reagent Plus (PerkinElmer Life Sciences). After stripping, membranes were incubated with anti-GAPDH antibody (Abcam, ab9485, 1/1000) and post-treated as above. Densitometry of the bands was quantitated with ImageJ.

### ASIC2 Immunohistochemistry

Pools of 15-20 rat CCDs were transferred to Superfrost Gold glass slides, fixed with 4% paraformaldehyde, incubated for 20 min at room temperature in 100 mM glycine, and permeabilized for 30 s with 0.1% Triton. After blocking, slides were incubated with anti-pan-ASIC antibody (Abcam, ab77384, 1/500) for 1 h at room temperature), rinsed and incubated 1 h at room temperature with and FITC-coupled anti-rabbit IgG (1/500), and observed on a confocal microscope (40, Zeiss observer.Z1, LSM710).

Two serial sections of human kidney biopsies were used for ASIC2 and aquaporin 2 (AQP2, a marker of the distal nephron) labeling respectively. ASIC labeling was realized as for isolated CCDs. For AQP2, after blocking, slides were incubated with and anti-AQP2 antibody (Santa Cruz, sc709882, 1/500) for 1 h at room temperature, rinsed and incubated 1 h at room temperature with TRITC-conjugated anti rabbit IgG (1/500).

### Albumin immunostaining

8μm rat kidney sections fixed with 4% paraformaldehyde and included in OCT were transferred to Superfrost Gold Glass slides, rinsed with PBS, incubated for 20 min at room temperature in 100mM glycine, permeabilized for 30s with 0.1% triton and incubated 5 min in SDS 1% for antigen retrieval. After blocking, slides were incubated with anti-AE1 antibody (gift from C. Wagner, Institute of Physiology, Zurich, Switzerland, 1/500) for 1h at room temperature, rinsed and incubated with TRITC-coupled anti-rabbit IgG (1/500) for 1h at RT. After rinsing, anti-albumin antibody FITC-coupled was added (DAKO F0117, 1/500).

### Quantification of phospho-ERK

5μm cryo-sections of rat kidney fixed with 4% paraformaldehyde were transferred to Superfrost Gold Glass slides, rinsed with PBS at 4°C and incubated for 10 min at 94°C in citrate buffer pH 6.0. After blocking with donkey serum (10%, for 30 min at room temperature) slides were incubated overnight at 4°C with anti-AQP2 antibody (Santa Cruz, sc-9882, 1/400) and anti-P-p44/42 MAPK (T202/Y204 antibody (Cell Signaling Technology, 02/2016) diluted at 1/400 each in PBS containing 5% milk and 0.01% Tween. After rinsing (3-time 5 min in PBS Tween 0.01%, 5min), slides were incubated for 1h at room temperature with donkey fluor 488-conjugated anti-goat IgG (Santa Cruz, sc-362255, 1/500) and donkey Alexa fluor 555-conjugated anti-rabbit IgG (Life Technology, 02/2016, 1/2000) and DAPI 1/2000. After rinsing and mounting in Dako Glycergel mounting medium, slides were observed on a confocal microscope (40, Zeiss observer.Z1, LSM710). Images were analyzed using ImageJ. The phospho-ERK fluorescence (red) in cross sections of AQP2-positive tubules (green) was measured and expressed as a function of the tissue area of the cross section. Fifteen-20 tubule sections were analyzed in each animal and the mean value was normalized to that of a same CC-control WT rat analyzed in each experiment.

### Statistics

Results are expressed as means ± SE from several animals. Comparison between groups was performed either by unpaired *t* test or by variance analysis followed by Fisher’s protected least significant difference test, as appropriate.

## Acknowledgements

A.S. and M.F. were supported by a grant from the Fondation pour la Recherche Médicale (grants #ING20121226213 & #FDM20091117400 respectively). This work was supported in part by a grant from the Fondation du Rein. BV was supported by the « Fonds pour la recherche Thérapeutique » (Lausanne), and by the SNF (Swiss National Science Fondation). Authors are grateful to Dr E. Lingueglia (IPMC, Valbonne, F-06560, France) for providing rat ASIC2a cDNA and for his helpful comments. Physiological analysis have been performed with the help of Gaelle Brideau and Nadia Frachon from the “platforme d’exploration fonctionnelle du petit animal” of the team “Physiologie Rénale et Tubulopathies” at the Centre de Recherche des Cordeliers (CRC). We are grateful for the technical assistance of the crews of the Centre d’Exploration Fonctionnelle (CEF) at the CRC in the management of our colony of rats.

## Authors’s contributions

AD, MF, AS, BV and GC designed the project. AD wrote the first draft of the paper. MF, LC, CW, MG and LC carried out animal experiments, biochemical, qPCR and immunolabeling analysis, MF, MP and GD carried out analysis of human samples, AS, GP, MK and NB carried out electrophysiological analysis, GB and LC performed cloning of ASIC2b variant. IA and SR generated the rat Asic2b KO and CW, GC and LC set up the colony of ASIC2b KO rats at the CEF animal facility and established the procedure for their genotyping. LM and PH carried out microperfusion on isolated tubules. All the authors discussed the results and commented the manuscript.

## Conflict of interest

The authors have declared that no conflict of interest exists.

**Supplementary Data. A.** *Accn1* gene (encoding for Asic2a and 2b proteins) structure and CRISPR/Cas9 targeting strategy. Single-guided RNA (sgRNA) was designed to target DNA sequence in exon 1 (flash symbol) specific of *Asic2b* and let *Asic2a* as a wild type sequence. **B.** *Asic2b* exon 1 (WT) with sequence targeted by the sgRNA (underlined plus PAM in bold) and the resulting deletion (Del). **C.** Example of gel electrophoresis showing the 20bp deletion in mutated rats compare to WT rats. **D.** Amino-acid sequence of Asic2b protein following nucleotide deletion by CRISPR-Cas9 process showing the position of a premature stop codon. **E.** Protein expression of Asic2b in nephrotic rat kidney homogenates from WT and Asic2b-null rats.

## Notes

### Competing Interest Statement

The authors have declared no competing interest.

