## Supplemental Figure 1 for "A new variant of ASIC2 mediates sodium retention in nephrotic syndrome"

Supplementary Data 1

A

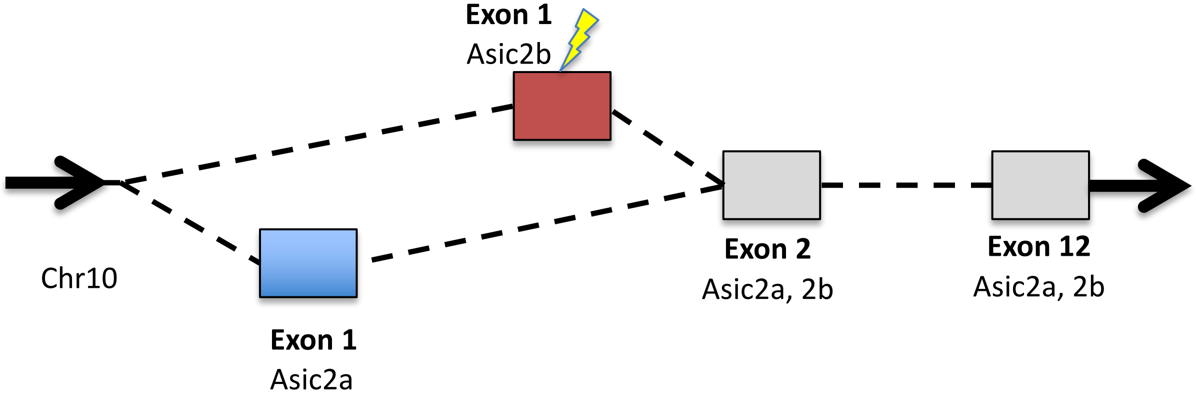


B

```
WT  CCGCAGGGGGCGGCCGTCCCTGAGTCGCACTAAATTGCACGGGCTGCGGCACATGTGCGCGGGGCGCACGGCGGCGGGAGGCTCTTTCCAGCGACGGGCGCTGT
Del CCGCAGGGGGCGGCCGTCCCTGAGTCGCACTAAATTGCACGGG--------------------GCGCACGGCGGCGGGAGGCTCTTTCCAGCGACGGGCGCTGT
```

C

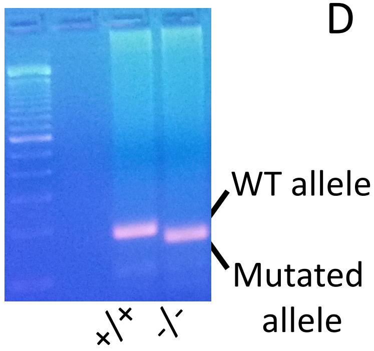


D

**Met** S R S G G A R L P A T A L S G P G R F R M A R E Q
P A P V A V A A A R Q P G G D R S G D P A L Q G P G V
A R R G R P S L S R T K L H G A H G G G R L F P A T G
A V G A G L L H V P R L A A V L V L E P P A L L A Q L
P V T H T S A P **Stop** V E P P A A V P R R H R V Q Q Q P
P A L P A P L Q G G P L L R G P L A R A A A S Q P H R
A P A G Q R A A A G R R A A P P V V P Q T G R L P P L
P A A A P L R G H Q R C L H G P F G P P A G G Y A A L
L Q V P G R A L W P A Q L L L ….

E

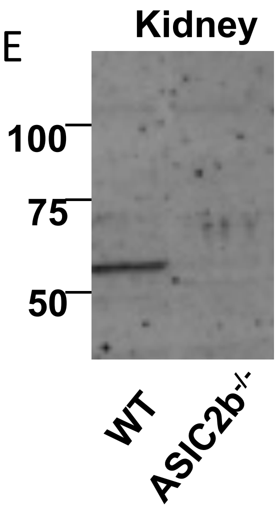
